# Microbial electrochemical systems for enhanced degradation of glyphosate: Electrochemical performance, degradation efficiency, and analysis of the anodic microbial community

**DOI:** 10.1101/2022.02.21.481054

**Authors:** Razieh Rafieenia, Mohamed Mahmoud, Fatma El-Gohary, Claudio Avignone Rossa

## Abstract

Glyphosate, one of the most used herbicides worldwide, is known as an aquatic contaminant of concern, and can present adverse impacts in agroecosystems. In this study, we investigated the degradation of glyphosate in microbial electrochemical systems (MESs), and analysed the microbial composition of enriched anodic biofilms, and comparing them with microbial communities of non-MESs enriched cultures. MESs supported higher glyphosate degradation (68.41 ± 1.21 % to 73.90 ± 0.79 %) compared to non-MESs cultures (48.88 ± 0.51 %). The Linear Sweep Voltammetry (LSV) analysis showed that MESs operated at +300 mV, produced a maximum current of 611.95 μA, which was the highest among all the applied voltages. 16S amplicon sequencing revealed a significant difference in microbial community composition of MESs anodic biofilms and non-MESs enriched communities. The anodic biofilms were dominated by *Rhodococcus* (51.26 %), *Pseudomonas* (10.77 %), and *Geobacter* (8.67 %) while in non-MESs cultures, methanogens including *Methanobrevibacter* (51.18 %), and *Methanobacterium* (10.32 %), were the dominant genera. The present study suggested that MESs could be considered as a promising system for glyphosate degradation.

## 1. Introduction

Pesticides are one of the most important contaminants of aquatic environments and present associated health risks to humans and animals, affecting diverse different ecosystems (Carneiro et al., 2015). Glyphosate (N-((phosphonomethyl) glycine, PMG) is a broad-spectrum herbicide, widely used for agricultural applications in more than 140 countries (Muñoz et al., 2021). The global use of glyphosate in 2014 reached 826 million kilograms, which showed a 12-fold increase compared to global usage in 1995 (Benbrook, 2016), and it is expected to reach 920 million kilograms by 2025 (Maggi et al., 2020).

The intensive use of glyphosate in the last decades has resulted in the contamination of surface and groundwaters through drainage, wind, soil erosion, and runoff (Samuel et al., 2017). The ecotoxicity of glyphosate has been reported to cause destruction in the balance of aquatic ecosystems (Muturi et al., 2017; Smedbol et al., 2018). Glyphosate concentrations in various aquatic environments have been reported to be in the range from ng. l^-1^ to 2.8 mg. l^-1^ (Lu et al., 2020). For instance, the highest residues of glyphosate in surface waters reached up to 700 μg. l^-1^ in Argentina (Peruzzo et al., 2008), 430 μg. l^-1^ in the USA (Mahler et al., 2017), and165 μg. l^-1^ in Europe (Villeneuve et al., 2011).

The presence of glyphosate in the environment is of concern, as it can adversely affect plants, animals, and human health and safety. Glyphosate is an endocrine disruptor in humans and might be carcinogenic in high concentrations (Andreotti et al., 2018). It has been reported that glyphosate can affect the composition of the gut microbiota and subsequently the central nervous system (Rueda-Ruzafa et al., 2019). In a recent study, the glyphosate concentrations in urine samples collected from 6848 French participants between 2018 and 2020, was analysed using ELISA. The results revealed that more than 99 % of the urine samples contained quantifiable levels of glyphosate (Grau et al., 2022). That study revealed that widespread contamination of glyphosate in France, an industrialized country, was mainly via ingestion, as individuals who consumed organic food and drank filtered water had lower levels of glyphosate in their urine. In terms of the effect on microorganisms, glyphosate exposure changed the composition of gut microbiota in rats, which was unexpectedly correlated negatively with male reproductive toxicity (Liu et al., 2022).

Several approaches can be used to treat glyphosate contaminated waters. Conventional methods including adsorption, and biological treatments, have been applied to treat water polluted with glyphosate (Villamar-Ayala et al., 2019). Also, advanced oxidation processes such as photocatalysis and electrochemical oxidation have been studied as alternative technologies for the removal of a variety of organic pollutants, including glyphosate (Feng et al., 2020). Biological treatments combined with physicochemical or advanced oxidation processes have been proposed as promising technologies for removal of a range of recalcitrant organic pollutants removal from wastewaters (Zhang et al., 2021). Among them, Microbial Electrochemical Systems (MESs), combining biological degradation with electrochemical oxidation, is considered as an emerging technology. MESs have been studied to remove a wide range of organic pollutants while recovering energy and valuable chemicals (Ramírez-Vargas et al., 2018; Wang and Ren, 2013). Due to the synergy between electrochemical processes and biological degradation, MESs could support higher removal of recalcitrant pollutants in comparison with biological degradation alone (Palanisamy et al., 2019). While microbial degradation of glyphosate has been studied in some detail (la Cecilia and Maggi, 2018), the use of microbial electrochemical technology for elimination of this pollutant has not been attempted so far.

In this work, we applied MESs for the degradation of glyphosate. We inoculated a natural microbial community in the anode of a glyphosate fed MES and studied the electrochemical performance of microbial populations and assessed the efficiency of the acclimatised biofilm to degrade glyphosate. The analysis of the composition of the microbial community allowed us to determine the abundance and diversity of the microbial species in the biofilms. The degradation of glyphosate and the composition of the microbial communities were compared with non-MESs cultures, which allowed us to find a correlation between the composition of the community and the enhanced degradation capacity. Finally, we assessed the electrochemical performance of enriched anodic biofilms of MESs.

## 2. Materials and Methods

### 2.1. Chemicals

All the chemicals including analytical grade glyphosate (Pestanal®) and potassium hexacyanoferrate (III) used for in the present study, were purchased from Sigma-Aldrich, UK. A stock solution of glyphosate (50 mg/L) was prepared dissolving the solid reagent in deionized water and stored at 4°C in dark conditions.

### 2.2. MESs configuration

Degradation of glyphosate was explored in two-chamber MES bioreactors operated in batch mode. The working volume of both cathode and anode compartments was 8 ml. Both anode and cathode electrodes (2.5 cm × 2.5 cm) were made up of carbon felt (Alfa Aesar, Haverhill, USA) and placed at equal distance from a cation exchange membrane (CMI-7000, Membranes Int., USA) separating the two chambers.

Duplicate batch experiments were performed at room temperature (25 ± 5°C) to study the degradation of glyphosate in MESs. Anaerobic digestion sludge collected from Goddards Green Wastewater Treatment Works (Hassocks, UK), was used to inoculate the anode compartment. The culture medium contained (per litre of deionized water): Na_2_HPO_4_ (6.02 g), KH_2_PO_4_ (1.024 g), NH_4_Cl (0.41 g), CH_3_COONa (1.0 g), and mineral media (10 mL). The composition of mineral media (per litre of deionized water) was CoCl_2_.6H_2_O (0.082 g), CaCl_2_.2H_2_O (0.114 g), H_3_BO_3_ (0.01 g), Na_2_MoO_4_.2H_2_O (0.02 g), Na_2_WO_4_.2H_2_O (0.01 g), MgCl_2_ (1.16 g), MnCl_2_.4H_2_O (0.59 g), ZnCl_2_ (0.05 g), CuSO_4_.5H_2_O (0.01 g), and AlK(SO_4_)_2_ (0.01 g) (Parameswaran et al., 2009). Nitrogen gas was sparged into the culture medium for 3 min to attain an anaerobic environment. The catholyte solution contained (per litre of deionized water) 4 g potassium hexacyanoferrate (III) and 100 ml PBS buffer and was replaced at the end of each feeding cycle to avoid acidification of the medium and its negative effect on current generation.

To enrich and acclimatize the anodic microbial community, MES bioreactors containing 10 μg/l glyphosate were operated initially as a microbial fuel cell with an external resistance of 1000 Ω, with an initial concentration of 10 μg/L glyphosate. The concentration of glyphosate was gradually increased to 500 μg/L with a weekly feeding regime lasting two months. Afterwards, the bioreactors were switched to microbial electrolysis cells to study the effect of applied voltage on glyphosate degradation. For this aim, an external resistance of 10 Ω was used while different voltages (0-500 mV) were applied to the bioreactors.

### 2.3. Analytical methods

At the end of each feeding cycle (one week), the anolyte solution was collected for analysis and the anode compartment was replenished with fresh medium. The anolyte samples were filtered using 0.2 μm membrane filters (CHROMAFIL® Xtra, Macherey-Nagel, Germany) and stored at -20°C for further analysis. Glyphosate concentrations were measured using a glyphosate ELISA kit (Abraxis, Eurofin Technologies, Hungary).

### 2.4. Electrochemical analysis

The electrical output of the MESs was recorded every 2 min using a data acquisition system (Pico Technology, Cambridgeshire, UK) connected to a personal computer. Current generation was calculated by Ohm’s law, I=V/R, where V is the reactor voltage and R is the external resistance (1000 or 10 Ω). Linear Sweep voltammetry (LSV) analysis with a scan rate of 5 mV/s, was carried out using a Palmsens potentiostat (PalmSens PS4, PalmSens, Houten, Netherlands) to study the activity of electroactive biofilms with and without applied voltages. Prior to LSV analysis, the anode chamber was filled with fresh medium, the catholyte solution was replaced, and the MESs were kept under open circuit conditions for 2 h.

### 2.5. Microbial community analysis

Total genomic DNA was extracted from the anodic biofilms, initial inoculum, and non-MESs cultures, using a DNeasy® PowerSoil® Pro Kit (Qiagen, Germany). To extract DNA from anodic biofilms, the MESs were disassembled at the end of operation, and the whole anode was used. The quality of DNA samples was analysed by Nanodrop (ThermoFisher Scientific), and 16S rDNA, 18S rDNA, and ITS amplicon sequencing was performed (Novogene (UK).

### 2.6. Phenotypic analysis

The enriched anodic community was studied using Biolog microplates. Biolog (MT2) 96-well microplates (Biolog, Hayward, USA) were used to assess the ability of initial inoculum and enriched anodic cultures to utilize glyphosate as a carbon source. Each well contains nutrient medium (except carbon source) and cellular respiration is measured spectrophotometrically by the reduction of the redox dye tetrazolium violet. In the present study, MT2 microplate assay was used to confirm if the microbial enrichment in MESs resulted in the enrichment of glyphosate-degrading microbes, which can use glyphosate as a sole carbon source. Microbial samples were extracted either from anodic biofilms or initial inoculum (sludge), washed three times and resuspended in sterile water. 15μl of a 2% glyphosate stock solution (0.3 mg of glyphosate) was added to each well. Resuspended inoculum or enriched microbial cultures (150 μl) were then inoculated into each well (triplicate). Microplates were incubated anaerobically at room temperature. The absorbance of the wells at different wavelengths (540, 565, 590, and 600 nm) was determined every 15 min using a Clariostar microplate reader (BMG LABTECH). The average well colour development (AWCD) curve was plotted after normalizing the absorbance of each well against the reading of the water-containing well. The negative values were considered zero (Koner et al., 2022).

## 3. Results and Discussion

### 3.1. Glyphosate degradation

Degradation of glyphosate by MESs and non-MESs enriched cultures was assessed using ELISA and the results are shown in Figure 1. In general, degradation of glyphosate in MESs reactors was significantly higher than in non-MESs reactors (68.41 ± 1.21 % to 73.90 ± 0.79 vs 48.88 ± 0.51 %, respectively). This difference in performance could be correlated to the differences in the composition of microbial communities between the two systems. The MESs showed higher abundances of *Pseudomonas* (10.77%), a genus that is known to be able to degrade glyphosate, compared to the abundance of *Pseudomonas* in non-MESs communities (2.11 %). In order to confirm that the enhanced glyphosate degradation in MESs was associated to the difference in microbial composition, we measured the glyphosate degradation in an abiotic electrochemical reactor (i.e., a MESs without inoculation). The comparatively low degradation (3.12 %) compared to that observed under biotic conditions, indicates that the contribution of abiotic degradation in MESs was insignificant compared to biodegradation.

**Figure 1.**
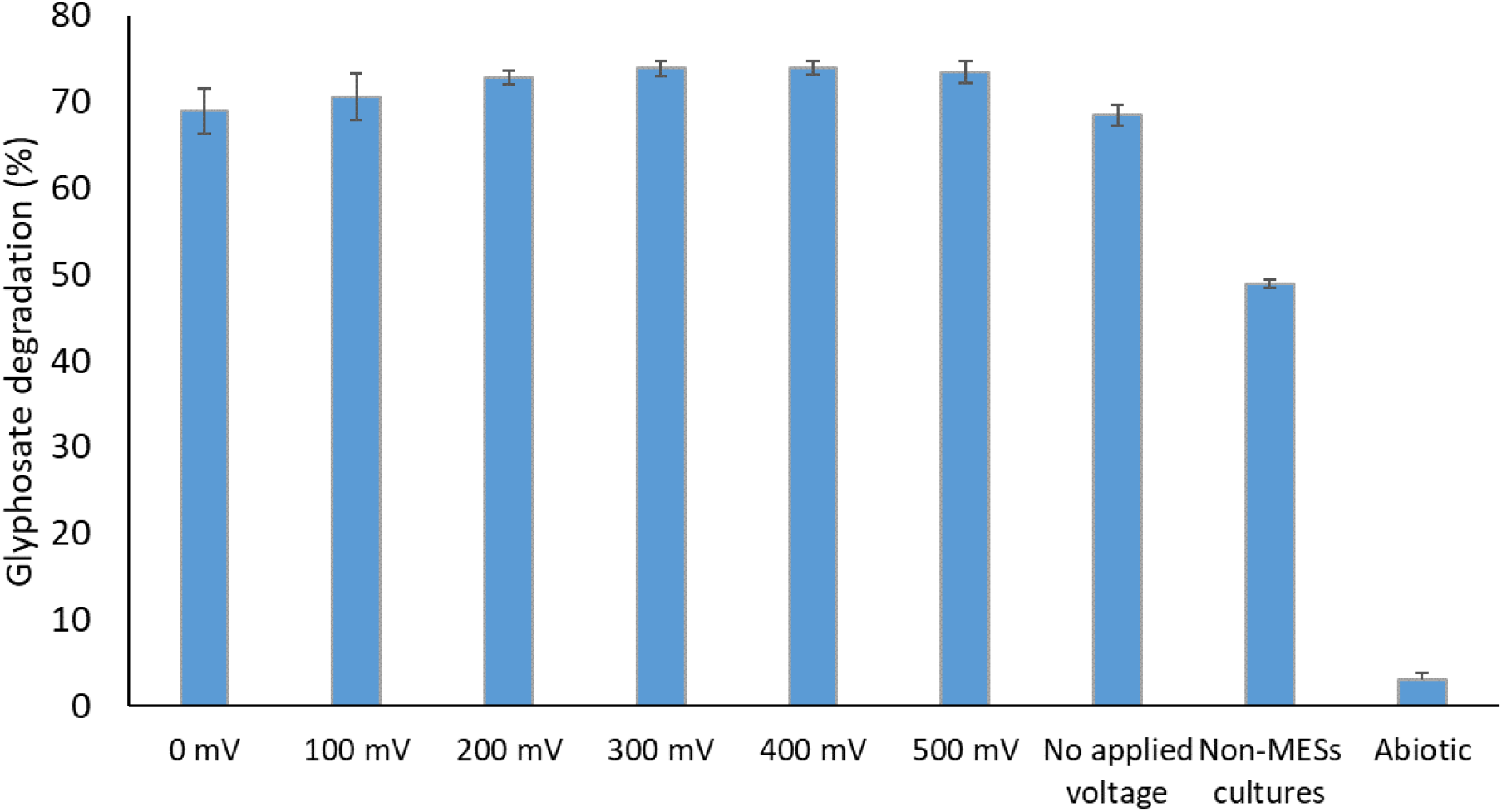
Degradation of glyphosate in MESs and non-MESs reactors

In general, glyphosate degradation in MESs with an applied voltage was slightly (2.09 – 5.49 %) higher than those without applied voltage. The degradation of glyphosate increased with applied voltage until reaching a value of 73.83 % at 300mV, remaining constant thereafter (73.90 % at 400 mV, and 73.39 % at 500 mV). It is known that optimum voltages can be determined for the microbial degradation of pollutants in MESs (Moghiseh et al., 2019). For example, an optimum voltage of 400 mV was determined for carbamazepine degradation in a MES. Higher voltages caused a decrease in carbamazepine degradation, a phenomenon attributed to the poor adaptability of anodic biofilm to high voltage (Tahir et al., 2019).

In order to assess the contribution of the electroactive microorganisms to the degradation of glyphosate, is necessary to compare the degree of degradation of the MES with that of a process in a non-MES reactor. Only a few studies on the biodegradation of glyphosate using microbial communities from aquatic or terrestrial environments have been reported. The half-life of glyphosate in most soils ranges between 7 and 60 days, depending on the microbial community and environmental conditions (Kanissery et al., 2019). In a study, the ability of microbial communities extracted from different soils for glyphosate biodegradation was evaluated and reported a 50% degradation of glyphosate (initial concentration of 50 μg/L) in 7 days (Forlani et al., 1999). In another study, half of the glyphosate was degraded in 9 days by downstream river microbial communities, dominated by *Phenylobacterium* sp. and *Starkeya* sp. (Artigas et al., 2020).

There are two major metabolic pathways for glyphosate biodegradation in glyphosate-degrading microorganisms. In microorganisms that can use glyphosate as a source of phosphorous, glyphosate is converted into sarcosine by the activity of enzymes able to cleave carbon–phosphorus (C–P) bonds. Sarcosine is subsequently degraded into carbon dioxide and water via different metabolic pathways. In microorganisms that use glyphosate as a nitrogen source, the metabolic pathway involves the conversion of glyphosate to aminomethylphosphonic acid (AMPA) and glyoxylate, by the activity of the enzyme glyphosate oxidoreductase, which splits the carboxymethylene–nitrogen (C–N) bond (Guijarro et al., 2018). According to the literature, most of the *Pseudomonas* species isolated as glyphosate-degraders metabolize glyphosate via sarcosine pathway (Singh et al., 2020).

The use of MESs for degradation of pesticides has been reported by a few studies, most of them using soil MFCs. The degradation of DDE (2,2-bis (p-chlorophenyl)-1,1-dichloroetylene), a persistent pesticide, in a soil MFCs operated for 2 months, 39.4 % of DDE was degraded in MFCs accompanied with a maximum power output of 30 μW (Borello et al., 2021). In another study, soil MFCs were used for remediation of hexachlorobenzene, an organic pesticide, and 37.12 % of the compound was degraded in 21 days (Cao et al., 2015). However, these cannot be compared to liquid MFCs.

### 3.2. Electrochemical characterisation of anodic biofilm

The electrochemical performance of the anodic biofilm was characterised using LSV to further confirm the enrichment of electroactive microorganisms and study the correlation between glyphosate degradation and electrochemical properties. LSV is a simple and widely used technique to study the electron transfer by anodic electroactive bacteria and subsequently the electrochemical performance of MESs. The polarisation curve indicates the relation between cell voltage and current generation for the studied voltage range and enables the calculation of the maximum current potential for an electrochemical system. Results of LSV analysis for MESs operated as MFCs and MESs with applied voltages are shown in Figure 2 and Figure 3, respectively. For the MFCs (Figure 2), the potential was swept linearly from open circuit voltage to 5 mV. The maximum current and power output obtained by MFCs were 196.11 μA and 60.29 mW respectively. An increase in the current was observed as the voltage swept from open circuit voltage to around 317 mV. No current increase was observed as the voltage swept beyond this point.

**Figure 2.**
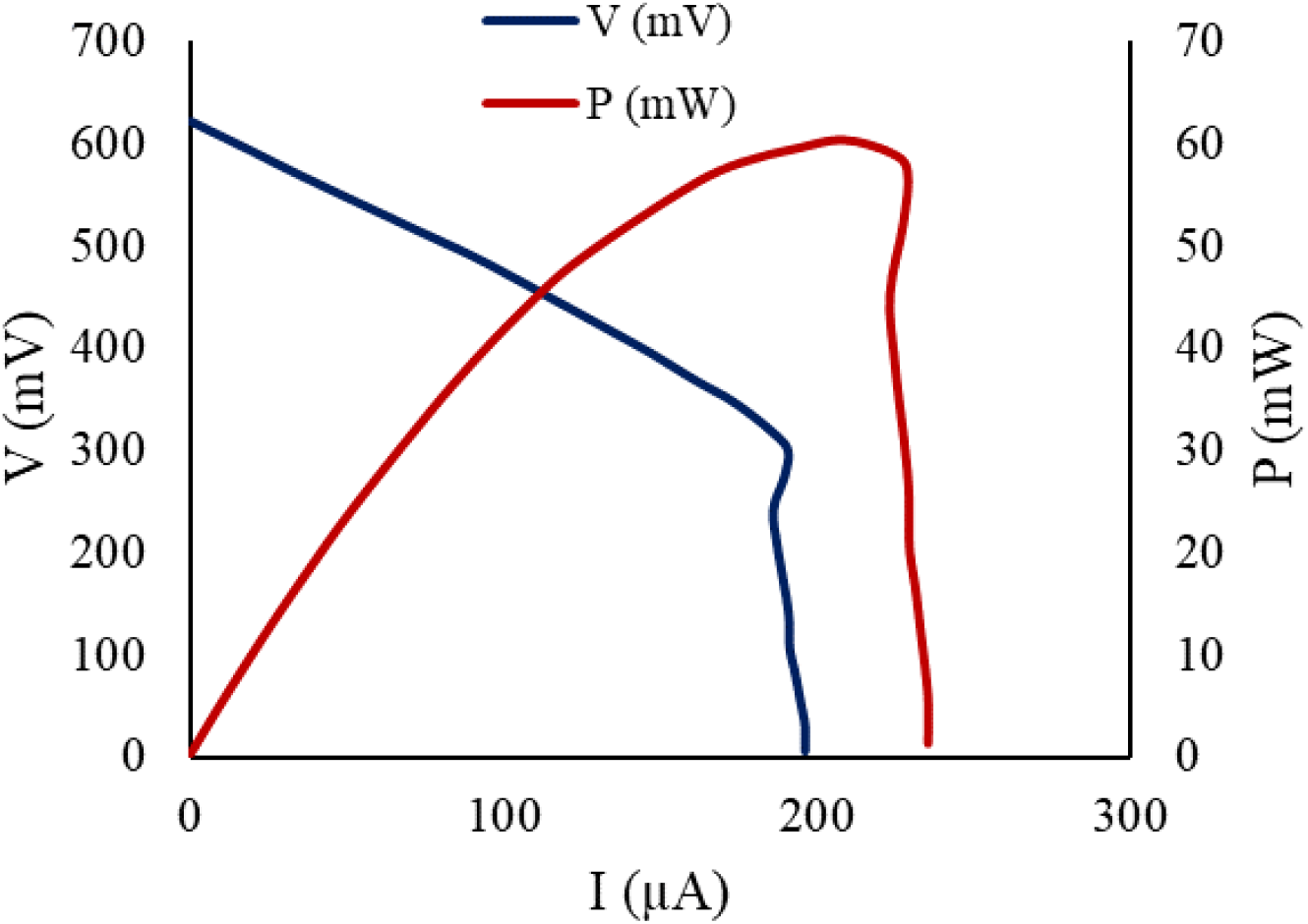
Polarisation curves and power outputs for the MFCs receiving 500 μg/L glyphosate

**Figure 3.**
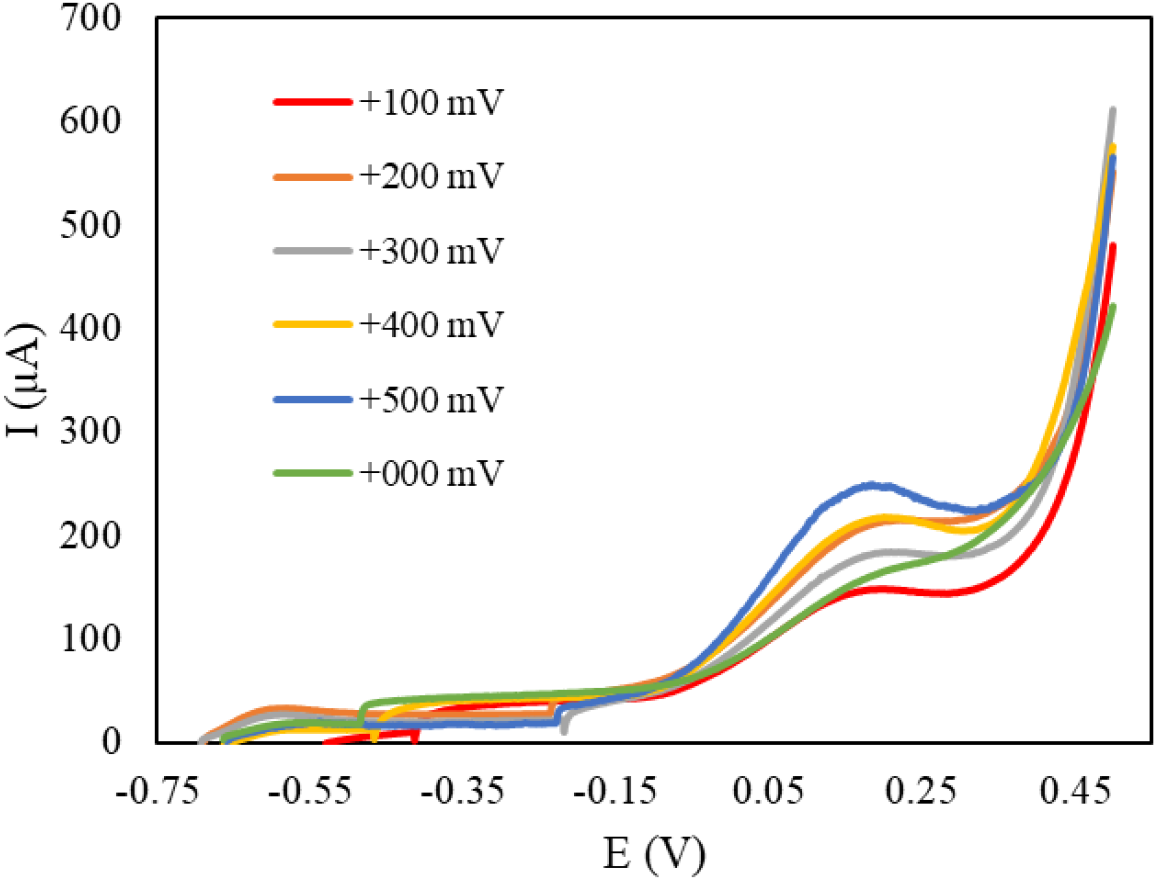
Linear Sweep Voltammetry analysis for the MESs with different applied voltages

LSV was also performed for the MESs operated with different applied voltages (from 0 to +500 mV), to study how anodic electroactive bacteria in each case respond to changes in cell voltage, and to determine the effect of the applied voltage on the electrochemical performance of MESs. To perform LSV analysis, the potential between the electrodes was swept linearly in time (starting from open-circuit voltage up to +500 mV) and the current response was plotted as a function of potential (Figure 3). An increase in the applied voltage from 0 to +300 mV led to an increased maximum current generation from 422.03 to 611.95 μA. This current increase could be attributed to the enhanced degradation of glyphosate, which resulted in excess electrons. Further increase in the applied voltage beyond +300 mV resulted in decreasing currents: the maximum currents for MESs operated with applied voltages of +400 and +500 mV were 575.68 and 565.44 μA, respectively. While the application of a potential stimulates EAB and therefore increase the current generation (Liu et al., 2015), voltages higher than a threshold value may reduce EAB activity due to changes in conformation, structure or redox properties of the enzymes involved, resulting in lower electrochemical performances (Dennis et al., 2016).

### 3.3. Microbial community analysis

The microbial communities of the initial sludge, the anodic biofilms, and the non-MES control cultures were analysed by 16S rRNA high-throughput sequencing to: 1) determine the changes in the composition of the enriched microbial populations caused by exposition to glyphosate; 2) to study the variation between microbial species enriched in MESs and non-MESs reactors; and 3) to gain further insight on the possible pathways for degradation of glyphosate in MESs.

The Goods coverage index for the initial sludge (0.993), MESs anodic biofilm (0.999), and non-MESs cultures (0.996), demonstrated that almost all the microbial populations and operational taxonomic units (OTUs) were covered by 16S rRNA sequencing.

The number of observed species, as well as the Chao1, Shannon and Simpson indices indicated that microbial diversity decreased after enrichment in either MESs or non-MESs reactors in comparison with the initial sludge (Table 1). Shannon diversity indexes of 3.531, 3.614, and 6.702, for the anodic microbial communities, the non-MESs cultures, and the initial inoculum, respectively, suggested that non-MESs enriched cultures were more diverse than MESs anodic biofilms. The lower diversities of enriched cultures could be attributed to the inhibition of glyphosate-sensitive species in both MESs and non-MESs reactors.

**Table 1.** Alpha diversity indices for different microbial samples

| Sample | Observed species | Shannon | Simpson | Chao1 | ACE | Goods coverage | PD_whole_tree |
| --- | --- | --- | --- | --- | --- | --- | --- |
| Initial inoculum | 2008 | 6.702 | 0.948 | 2278.905 | 2340.709 | 0.993 | 167.699 |
| MES | 428 | 3.531 | 0.722 | 458.373 | 475.508 | 0.999 | 39.000 |
| Non-MES | 703 | 3.614 | 0.720 | 784.657 | 830.485 | 0.996 | 81.674 |

The results of microbial community analysis at phylum, class, and genus level are shown in Figure 4. The initial community (inoculum) was dominated by five phyla, including *Euryarchaeota* (27.01%), *Proteobacteria* (16.75%), *Firmicutes* (14.86%), *Cloacimonetes* (14.03%), and *Bacteriodetes* (10.18%). After the enrichment process, there was a huge difference between microbial communities detected in anodic biofilms and non-MESs cultures. In both MESs and non-MESs enriched cultures, some phyla including *Firmicutes, Cloacimonetes, Bacteriodetes, Chloroflexi*, and *Spirochaetes*, exhibited an obvious decrease compared to the initial inoculum, attributable to their sensitivity to glyphosate. *Actinobacteria* (51.74%) was the dominant phylum in anodic biofilms, while in non-MESs cultures, *Euryarchaeota* showed the highest relative abundance (68.06%). *Proteobacteria* was the second dominant phylum in both anodic biofilms (31.18%) and non-MES cultures (17.73%) (Figure 4a). The selective enrichment of EAB in anodic biofilms was further confirmed at class level with increased relative abundances of *Actinobacteria* and *Gammaproteobacteria* (about 12.8 and 2-fold increases compared to the initial sludge, respectively) (Figure 4b). *Actinobacteria* (51.68 %), *Gammaproteobacteria* (18.67 %), and *Desulfuromonadia* (8.67 %), were the three classes with highest relative abundances in anodic biofilms, while in non-MESs enriched cultures, *Methanobacteria* were predominant (63.2 %). Some classes including *Clostridia, Bacteroidia, Alphaproteobacteria*, and *Anaerolineae*, showed lower abundances in both MESs and non-MESs cultures, compared to the inoculum.

**Figure 4.**
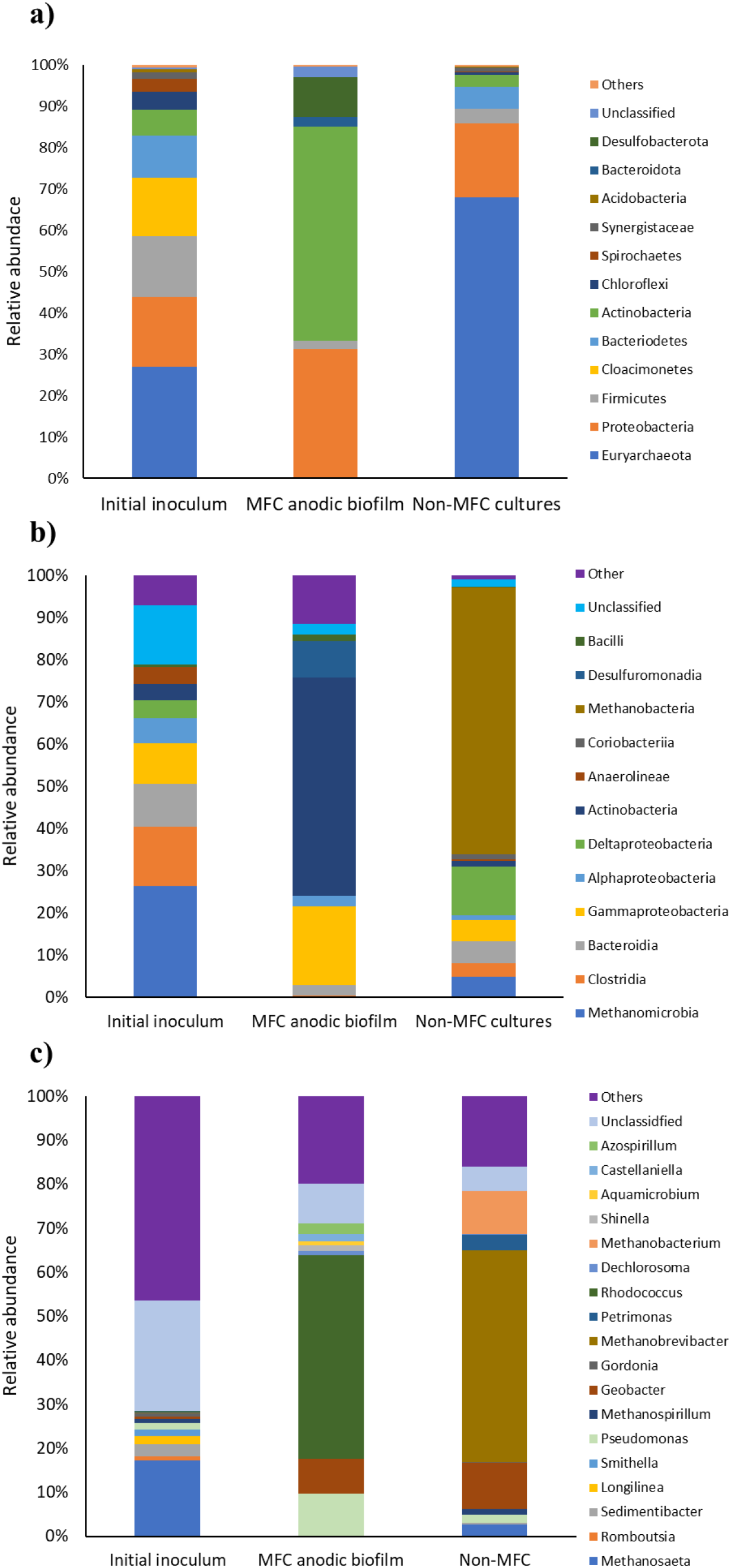
Composition of microbial communities on electrodes and anaerobic batch cultures at a) phylum, b) class and c) genus level

Genus level analysis (Figure 4c) indicated that the anodic biofilm was dominated by three genera known for their electroactivity, including *Rhodococcus* (51.26 %), *Pseudomonas* (10.77 %), and *Geobacter* (8.67 %), while *Methanobrevibacter* (51.18 %), *Geobacter* (11.20 %), and *Methanobacterium* (10.32 %) had the highest relative abundances in non-MESs cultures. Interestingly, methanogens including *Methanosaeta, Methanobacterium*, and *Methanobrevibacter* were not detected in anodic biofilms, suggesting that EAB outcompeted methanogens. On the contrary, the three mentioned methanogenic genera were enriched significantly in non-MESs cultures, together accounting for more than 60% of community. Members of the genera *Rhodococcus, Geobacter*, and *Pseudomonas*, are widespread in different environments including soil and marine sediments, and can consume a wide range of recalcitrant organic compounds as substrate (Yano et al., 2014). In addition, members of these three genera have been detected frequently in MESs (Allam et al., 2021; Shen et al., 2020). Several of the most common glyphosate-degrading species belong to *Pseudomonas* genus (Zhan et al., 2018). It has been reported that the abundance of *Pseudomonas* to be related with degradation of glyphosate in soils (Gimsing et al., 2004).

Unclassified genera accounting for 10.11 % and 5.90% in anodic biofilms and non-MESs cultures respectively, may also have important roles in glyphosate degradation. Other identified genera that have currently been isolated as glyphosate-degraders include *Achromobacter, Agrobacterium, Comamonas, Achromobacter, Ochrobactrum, Geobacillus* and *Rhizobium* (Massot et al., 2021; Zhan et al., 2018). In the present study, none of these genera were found in the initial inoculum nor after microbial enrichment.

Our shows that some of the electrogenic bacteria including *Rhodococcus, Geobacter*, and *Pseudomonas*, could tolerate glyphosate and be enriched in MESs. These genera have been previously reported to participate in the biodegradation of other pollutants including antibiotics, herbicides, pharmaceuticals, and aromatic compounds (Hou et al., 2020; Xie et al., 2021; Zhu et al., 2021).

### 3.4. MT2 microplate assay for glyphosate utilization by anodic communities

MT2 assay is an inexpensive and effective method to evaluate the ability of inoculated microorganisms to utilize selected carbon sources. An MT2 microplate assay was carried out to further study the bioactivity of anodic biofilm for glyphosate utilization. The AWCD values of the glyphosate utilized by anodic microbial community was significantly higher than those of the initial inoculum (Figure 5). This suggest that the microbial enrichment in MESs resulted in enhanced metabolic activities for glyphosate utilization. The complementary results obtained from MT2 microplate assay, together with glyphosate degradation values obtained by ELISA, confirmed that MESs operation resulted in enrichment of microbes that are involved in glyphosate degradation. Moreover, MT2 microplate assay revealed that anodic microbial communities could grow on glyphosate as a sole carbon source.

**Figure 5.**
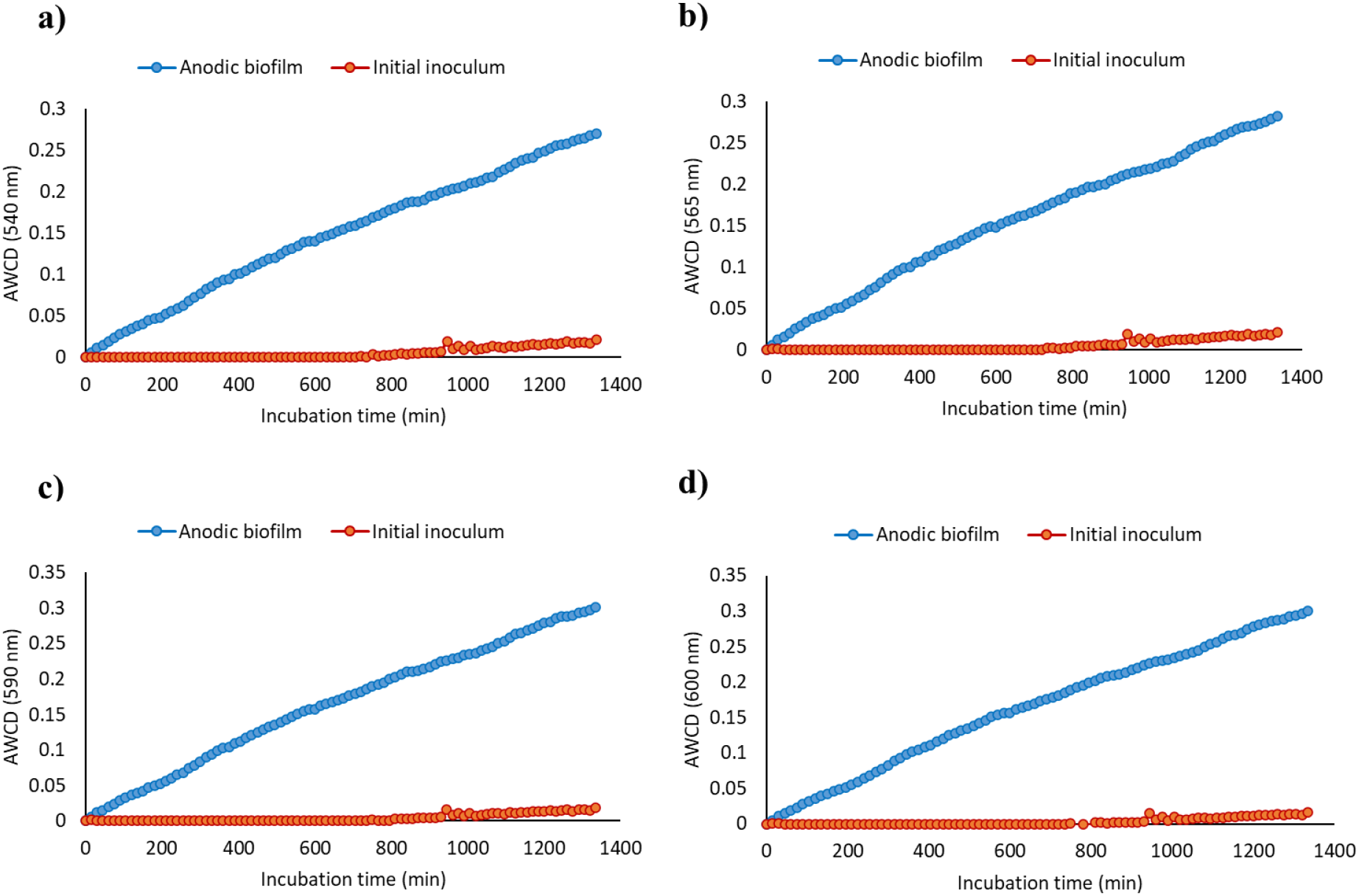
Average well colour development (AWCD) at different wavelengths for glyphosate utilization by initial inoculum and enriched anodic biofilm

## 4. Conclusions

The present study reported for the first time the use of MESs for the enhanced degradation of glyphosate. The preliminary results suggested than MESs could support higher degradation of glyphosate (68.41 ± 1.21 % to 73.90 ± 0.79 %) compared to biological degradation (48.88 ± 0.51 %). Microbial community analysis revealed that abundant genera in anodic biofilms of MESs were *Rhodococcus, Pseudomonas*, and *Geobacter*, while the microbial community enriched in non-MESs reactors, was dominated by methanogens (*Methanobrevibacter* and *Methanobacterium*). The significantly low degradation of glyphosate in abiotic conditions (3.12 %), revealed the enhanced degradation of glyphosate in MESs could be attributed to the difference in microbial community composition.

Our study shows the enhanced degradation of glyphosate using MESs. To further assess the fate of glyphosate and identify or confirm its biodegradation pathways by a microbial community, and the mechanisms involved, the expression levels of the genes related to different degradation pathways could be monitored. In addition, isotope labelling could be used to trace its biodegradation metabolites.

The combination of MES with other advanced oxidation processes could result in the complete elimination of glyphosate from polluted wastewaters.

## Acknowledgments

RR, MM, FE-G and CAR are supported by an Institutional Link Grant from the British Council (Newton-Mosharafa Fund) (Grant no. 352368074) and the Science, Technology, and Innovation Funding Authority (STIFA), Egypt (Grant no. 30901). MS and CAR would like to acknowledge support by the EU Horizon 2020 Research project GREENER (Grant Agreement No. 826312).

